# A Cartography of Differential Gene Methylation in Myalgic Encephalomyelitis/Chronic Fatigue Syndrome: Different Network Roles in the Protein-Protein Interactions Network Play Different, Biologically Relevant, Roles

**DOI:** 10.1101/2021.12.20.473375

**Authors:** Aurelia Wilberforce, Giulio Valentino Dalla Riva

**Author notes:** **Correspondance to:** Aurelia Wilberforce —.

## Abstract

Myalgic Encephalomyelitis, or Chronic Fatigue Syndrome (ME/CFS), is characterised by severe fatigue and associated with immune dysfunction. Previous studies of DNA methylation have found evidence of changes in immune cells for ME/CFS. However these studies have been limited by their small sample size. Here, we aggregate three comparable datasets to achieve a larger sample size and detect small changes to DNA methylation. We find 10,824 differentially methylated genes, with a small average change. Next, from the currently known interactions of relevant proteins, we build a Protein-Protein interaction network and,localising the network cartography analysis, we identify 184 hub genes. We find that different hub types play different, and meaningful, biological roles. Finally, we perform Gene ontology enrichment analysis, and we find that these hubs are involved in immune system processes, including response to TGF-*β* and LPS, as well as mitochondrial functioning, supporting previous theories about ME/CFS. We also show that dopaminergic signalling may potentially contribute to immune pathology in ME/CFS, suggesting a possible interplay with Long Covid. Our results demonstrate the potentiality of network analysis in shedding light on the epigenetic contribution to the immune dysregulation of ME/CFS.

## 1 Introduction

Chronic Fatigue Syndrome, also known as Myalgic Encephalomyelitis (ME/CFS), is a devastating and heterogeneous disease characterised by severe fatigue that is made worse by exertion (Carruthers et al., 2011; Fukuda et al., 1994). The disease may result from immune dysregulation (Cortes Rivera et al., 2019; Komaroff, 2019; Missailidis et al., 2019; Morris and Maes, 2013); in some cases, in response to previous viral infection, such as Sars-Cov-2 (Covid-19) (Komaroff et al., 2021). The symptoms of Long Covid are similar to ME/CFS, but it is too early to fully delineate the relationship between these two illnesses (Jason et al., 2021; Wong and Weitzer, 2021). It is important to understand the pathophysiology of ME/CFS, and its potential implications on Long Covid.

Despite the huge impact ME/CFS has on patients, efforts to find stable immune abnormalities have been largely unsuccessful. This is especially true of the investigations of cytokines, which often have conflicting results (Blundell et al., 2015; Corbitt et al., 2019)—cytokines function in an autocrine and paracrine manner, so their levels in the blood do not necessarily represent levels of inflammation elsewhere (Vanelzakker et al., 2019). Different studies, such as (Ekua Weba Brenu et al., 2014; Cliff et al., 2019; Curriu et al., 2013), have identified varying perturbations to the T cell populations, B cells, and natural killer (NK) cells. The source of these varying changes is unclear. Still, it is likely that the pathophysiology of ME/CFS is dominated by the effects of this immune activation and inflammation (Cortes Rivera et al., 2019; Komaroff, 2017; Morris and Maes, 2013).

Attempts at finding a genetic origin, and in particular epigenetic changes in immune cells of ME/CFS, have been partially successful (E. Brenu et al., 2014; De Vega et al., 2017, 2014; Helliwell et al., 2020; Herrera et al., 2018; Trivedi et al., 2018; Vega et al., 2018). DNA methylation is an epigenetic modification to DNA strands which can affect gene transcription (Moore et al., 2013). Methylation can be inherited but can also be modulated by environmental circumstances during a person’s life (Martin and Fry, 2018; Matosin et al., 2017). As a result, if there are methylation changes in ME/CFS immune cells, these may function differently to that of healthy controls, even in response to habitual levels of signalling molecules (such as cytokines). Previous studies have been often limited by small patient samples, precluding the detection of small changes in gene methylation. Data acquisition through clinical procedure is a delicate, time and resource consuming, task. Yet, the volume of publicly available datasets is rapidly growing.

Here, we hypothesise the presence, in the cells of patients with ME/CFS, of a large number of slightly differentially methylated genes (DMGs) potentially missed by studies with smaller sample sizes. To test this, we firstly merged and reconciled three highly comparable studies with publicly available data (De Vega et al., 2017, 2014; Herrera et al., 2018). Secondly, we built a Protein-Protein interaction (PPI) network, looking for known interactions of proteins corresponding to differentially methylated genes in the STRING database (Szklarczyk et al., 2021, 2019). Then, adapting the notion of network cartography (Guimerà and Amaral, 2005) to our scenario, we identified network genes playing a key role between the DMGs (which we call *hubs*). Finally, we conducted a Gene Ontology (GO) enrichment analysis of these hub genes.

Our data enrichment and network analysis shed light on the origins of ME/CFS immune pathology. In particular, we found that the hub genes, 0.017% of the differentially methylated genes, interact with over half of the PPI network. Moreover, the different hub types play distinct, and biologically relevant, roles in ME/CFS pathology as shown both by their network properties and by the results of the GO enrichment analysis. Overall, we found that terms associated to response to cytokine, lipopolysaccharide (LPS) and transforming growth factor beta (TGF*β*) feature in the GO analysis, as well as terms related to dopaminergic processes. This suggests that dopaminergic deregulation may underlie some immune dysfunction in ME/CFS. It is not yet clear whether long covid will share dysregulated responses to these molecules, however dopaminergic processes may play a role in long covid as dopamine has already been shown to play a role in COVID-19 itself Nataf (2020a).

## 2 Materials and methods

In order to shed light into the epigenetic contribution to the immune dysregulation of ME/CFS, we applied a four stepped analytic workflow (fig. 1), described in more details in the following sections.

**Figure 1.**
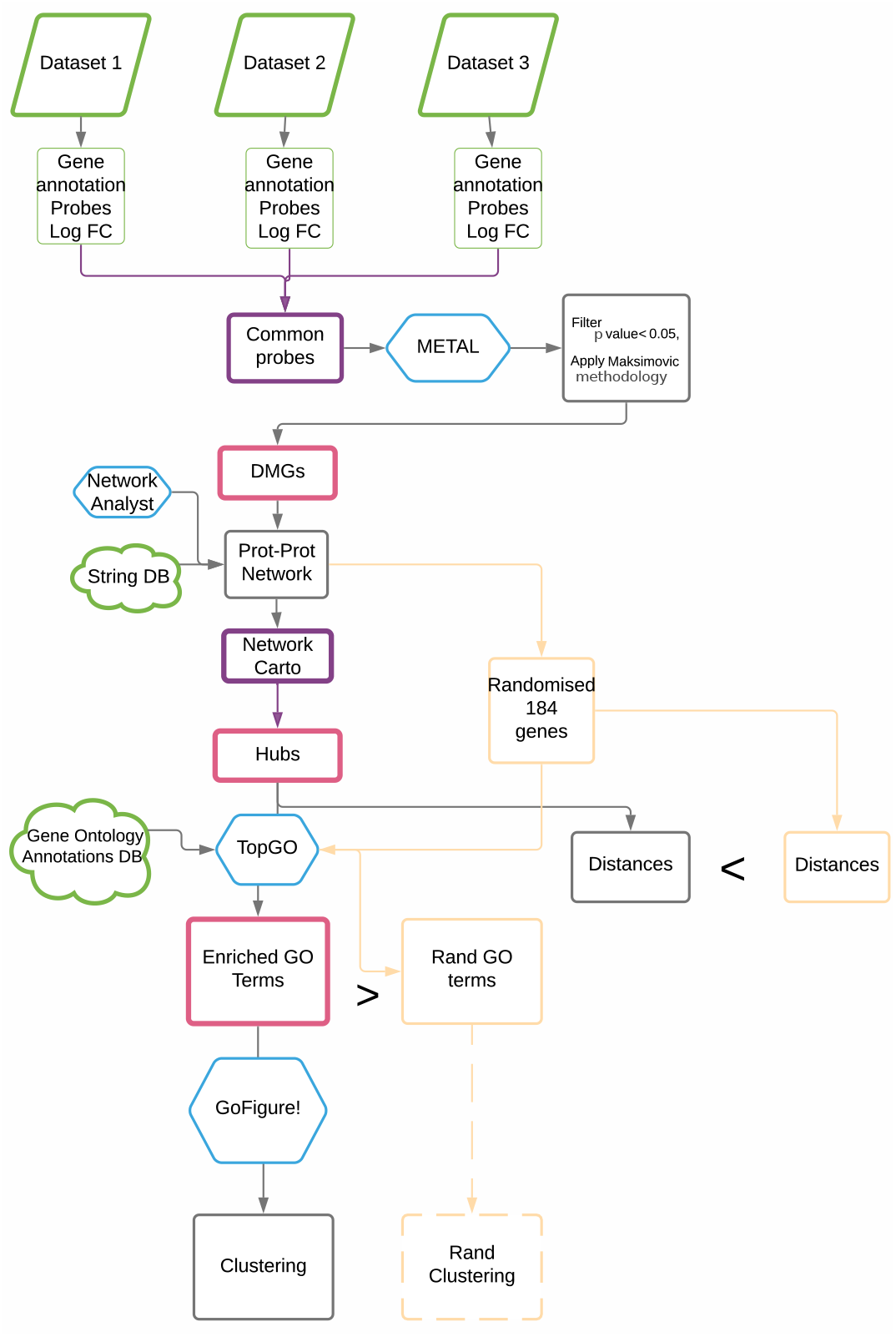
A schematic view of our analytical pipeline. In green the external data sources we used in our analysis. In purple the novel steps we introduce, namely: the merging of three different datasets and the localisation of Guimerà and Amaral (2005) network analytic framework. In pink the outcomes of our analysis (a list of differential methylated genes, a classification of genes in differen network hub types, and a list of enriched Gene Ontology networks). In azure some of the software we used. In yellow the analytic pipeline for the validation (random) data: as explained in the results section, we could not perform clustering on the validation GO terms because not enough terms we significant.

Shortly, we first merged three different, but comparable, datasets (De Vega et al., 2017, 2014; Herrera et al., 2018), from which we determined Differentially Methylated Genes (DMGs) (Maksimovic et al., 2021) in ME/CFS. Next, starting from the proteins associated to those DMGs, we built a PPI network assembling PPI known in humans. Then, localising Guimerà and Amaral (2005)’s network cartography, we identified key genes (*hubs*) and characterised their role in the PPI network (hub *types*). Finally, we determined enriched Gene Ontology terms associated to the hubs and identified themes.

We validated our results contrasting them with those of random samples of a adequately sized sets of genes.

### 2.1. Data

From NCBI’s Gene Expression Omnibus website (GEO, (Barrett et al., 2012; Edgar et al., 2002)), we obtained three datasets covering gene methylation of peripheral blood mononuclear cells (PBMCs) in ME/CFS (De Vega et al., 2017, 2014; Herrera et al., 2018) (accession numbers GSE93266, GSE59489, and GSE156792 respectively). All three datasets were produced using the platform GPL13534, Illumina HumanMethylation450 BeadChip. The data is analysed using the GEO2R service (Barrett et al., 2012) provided by GEO. The processed data provided by the authors of the original studies is analysed by GEO2R and each probe is given a log fold change (LogFC) and p-value. We performed p-value adjustment via false discovery rate (FDR) as suggested by Benjamini and Hochberg (1995).

We identified the probes that were assessed in all three studies. Each probe, for each study, was annotated with an effect direction based on the sign of log fold change (LogFC) and given a weight based on the sample size of the relevant study. We imported probe identifier, adjusted p-values, effect direction and study weight in METAL (Willer et al., 2010) and performed a sample size based analysis. We filtered for probes with a p-value less than 0.05, which we annotated with associated gene names and gene regions as found in the original datasets. We discarded probes not related to a gene or associated to a contradictory effect direction across studies. Analysis were performed in R (Team, 2021).

### 2.2. Probe annotation and gene expression

Starting from the identified significant probes, we proceeded to identify genes of interest for six gene regions: TSS200, TSS1500, Body, 3’UTR, 5’UTR, and 1st exon. We considered for analysis all the genes present in probe with a splice variant corresponding to one of the six regions.

We approximated gene methylation level in each region covered by our study following the work of Maksimovic et al. (2021). Methylation in different gene regions results in differing effects on expression (Moore et al., 2013; Varley et al., 2013). The most consistent relationship is an inverse relationship between promoter associated regions and gene expression (Martino and Saffery, 2015; Moore et al., 2013). This may also apply to the 5’UTR and 1st exon regions (Brenet et al., 2011; Moore et al., 2013; Varley et al., 2013). We shall call these regions the ‘inverse regions.’ In the gene body and 3’UTR, the relationship with gene expression is more complex but may be positive (Martino and Saffery, 2015; Maussion et al., 2014; McGuire et al., 2019; Yang et al., 2014). These we denote the ‘positive regions’ We calculated average LogFC for each analysed probe as the mean of the LogFCs over each study per probe. Let *G*_*R*_ be the total (weighted) LogFC of a gene *G* over a certain region *R*, that is, the sum of the (weighted) LogFCs over all the probes’ in region *R* annotated to that gene. For the inverse regions, *G*_*R*_ < 0 indicates upregulated gene expression; for the positive regions *G*_*R*_ > 0 indicates upregulated gene expression. Therefore, we classified genes with *G*_*R*_ greater than zero as potentially having upregulated expressiion and genes with *G*_*R*_ less than zero as potentially having down regulated expression. For potentially upregulated genes, we added the sum of *G*_*R*_ in inverse regions per gene and subtracted the sum of the *G*_*R*_ of the positive regions; gene regions with a *G*_*R*_ suggesting down regulation would cancel out a degree of the *G*_*R*_ suggesting upregulation. For potentially upregulated genes we defined their total LogFC as *G*_*t*_ = ∑_⊕_ *G*_*R*_ − ∑_⊖_ *G*_*R*_ where ⊖ and ⊕ denote the inverse and positive regions respectively. Then, we discarded any upregulated gene with a negative total logFC, *G*_*t*_ < 0. Similarly, for potentially downregulated genes we defined their total LogFC as *G*_*t*_ = ∑_⊖_ *G*_*R*_ − ∑_⊕_ *G*_*R*_ and we discarded any gene with a positive total logFC, *G*_*t*_ > 0. More details in the Supplementary Material.

### 2.3. Protein-Protein interaction network

We built a PPI network from potentially upregulated and potentially downregulated genes, using Network Analyst (Zhou et al., 2019) and the STRING database (Szklarczyk et al., 2021, 2019). The network was constructed such that each node represents a gene, and an edge connecting two nodes represents a known interactions between the proteins encoded by the genes. The PPI came from the STRING database and required a confidence score of at least 900 (denoted ‘highest confidence’ by (Szklarczyk et al., 2019)). The confidence score is calculated by STRING and based on the quality of each source of evidence for an interaction (Von Mering, 2004). The only genes considered for the network are the differentially methylated genes previously identified. From the DMGs, we built a zero-order network and extracted for analysis its giant connected component, which represents the PPI network in our study.

Our method for identifying hubs builds upon the notion of network cartography, introduced by Guimerà and Amaral (2005) and employes two indices. First, we defined the normalised degree, *z*_*i*_, of a node *i* in the the network as its scaled and centralised degree: 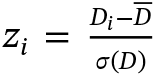, where *D*_*i*_ is the degree of the node *i*, and 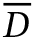 and *σ*(*D*) are the the average and the standard deviation of the degrees of all nodes in the network respectively. Next, let ⋆(*i*) be the induced subnetwork containing *i* and all the nodes at most one step away from *i*, see fig. 2 and let 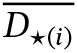 be the average node degree over ⋆(*i*). We defined the local participation coefficient of *i*, *P*_*i*_, as 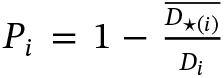. The index *P_i_* quantifies the connectivity of a node *i* to its local community. The original method by Guimerà and Amaral (2005) would require the identification of a community structure (clustering) in the PPI network to compute the participation coefficient; here, due to the constraints imposed by the sparsity of the network reconstruction and the lack of an established methodology to verify the communities, we relied instead on an indices defined either globally, *z*_*i*_, or locally over the neighbourhood of each node, *P*_*i*_. A direct comparison between local and original indices is offered in the Supplementary Material.

**Figure 2.**
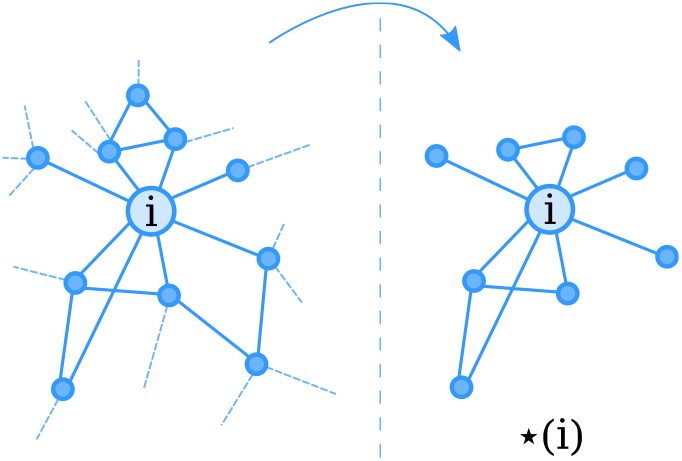
For any node *i* in our network, we denote ⋆(*i*) the network induced by the first-order neighbours of *i*. That is, the set of nodes of the network ⋆(*i*) is given by *i* and the all nodes at most one step away from *i*; the edges are given by all the edges in the original network between any two node in the set of nodes of ⋆(*i*). The figure shows which nodes and edges from the original network (left) are preserved in the induced network (right).

Using the normalised degree and the participation coefficient of each node, each gene was assigned a role based on the cutoffs suggested by Guimerà and Amaral (2005). Specifically, nodes with a normalised degree score *z*_*i*_ ≥ 2.5 were designated as hubs and assigned to three hub types based on participation coefficient. We named them

- **city hubs** (*P*_*i*_ ≤ 0.3): their neighbours area densely connected;
- **suburban hubs** (0.3 < *P*_*i*_ ≤ 0.75): an intermediate role, bridging between sparsely connected cliques of densely connected neighbours;
- **transit hubs** (0.75 < *P*_*i*_): their neighbours are sparsely connected.

Finally, we assessed the centrality of the hubs in terms of distances. As we are interested in the behaviour of the hubs as a set, and not individually, for each node in the network we computed its minimum distance to any of nodes in the set of all hubs (and any of the nodes in the sets of city hubs, in the set of suburban hubs, and in the set of transit hubs separetely).

We computed network indices and distance distributions using Graphs (James Fairbanks and Karpinski, 2021) in Julia (Bezanson et al., 2017). Code availalbe in the Supplementary Material.

### 2.4. Gene Ontology enrichment analysis

We performed gene ontology (GO) enrichment analysis using the bioconductor package topGO (Alexa et al., 2006; Alexa, 2021) in R (Team, 2021). We used a list of the hub genes abnormally methylated in ME/CFS, with a background list of all genes assayed by the 450k array so to identify significantly enriched GO terms. We used the Gene Ontology Biological Processes (GObp) dataset (Ashburner et al., 2000; “The gene ontology resource: Enriching a GOld mine,” 2021) and an elimination algorithm provided by topGO (Alexa et al., 2006; Alexa, 2021; Ashburner et al., 2000; “The gene ontology resource: Enriching a GOld mine,” 2021), with a threshold of at least 3 genes being annotated to a GObp term for inclusion in the analysis. Raw p-values provided by topGO were corrected for false discovery rate. With the same methodology, we analysed the genes of each hub type separately.

Using GO-Figure! (Reijnders and Waterhouse, 2021), we visualised the statistically significant GO terms from all hubs together as well as each hub type separatelly; assessed semantic similarity; and identified clusters in the semantic space, considering a similarity index cutoff (*si*) of *si* = 0.45, *si* = 0.5, and *si* = 0.7. GO-Figure! (Reijnders and Waterhouse, 2021) assigned cluster names based on a list of terms manually selected from the most general terms for each possible process within the full list of GO Terms (see Supplementary Material).

### 2.5. Validation

To validate our results, we compared them to those we would get by sampling at random equally sized set of the genes from the DMGs. For each random sample we repeated the same analysis as above. We ran a total of 200 repetitions for the GO enrichment and 1000 repetions for the distances distribution analysis.

#### 2.5.1. Reproducibility

Elaborated data and scripts can be found in a public repository github.com/officinadata/CFS_Proteomic.

## 3 Results

### 3.1. Probes and genes

Combining the three datasets, we obtained a total of 478,165 probes to analyse. Filtering for statistical significance at a level of *α* = 0.05, we reduced the set to 42,439 probes, of which 32,318 associated with a gene. We computed total methylation change and looked for evidence of up or down regulation, finding 10,824 genes differentially methylated—5,532 genes up regulated and 5,292 down regulate (more details in tbl. 1). Next, we built a zero-order network from the 6,470 genes with a sufficiently large absolute average LogFC, |*G*_*s*_| > 0.01.

**Table 1.**
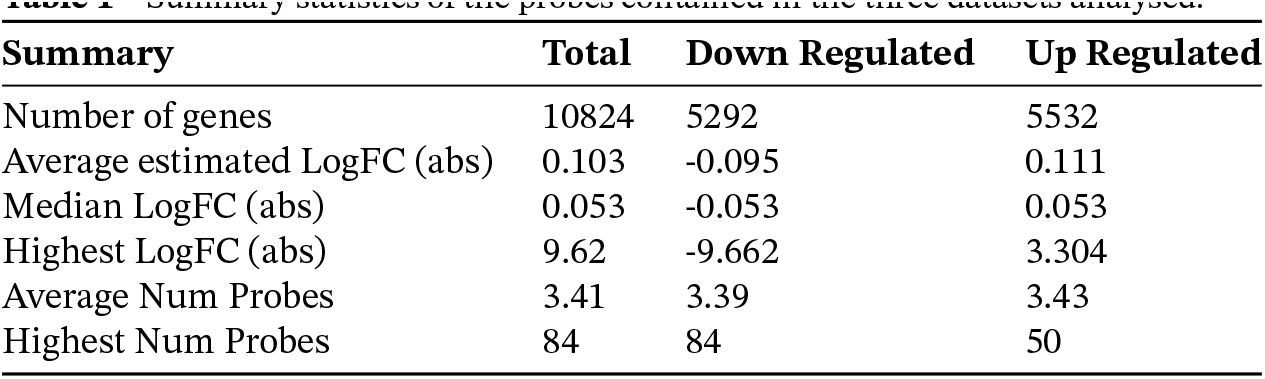
Summary statistics of the probes contained in the three datasets analysed.

### 3.2. Hub Identification

From the zero-order network, we extracted for analysis its giant component, which contained 4,770 nodes (coded by genes) and 40811 edges (interactions between two proteins). We computed *z*_*i*_ and *P*_*i*_ scores for each node in the PPI network, and filtered as hubs those 184 genes with a *z*_*i*_ ≥ 2.5. We found that 63 of the associated genes to be potentially downregulated and 121 potentially upregulated. We classified hubs based on their *P*_*i*_ scores into 61 Transit hubs, 40 suburban hubs, and 83 City hubs, than can be seen in fig. 3

**Figure 3.**
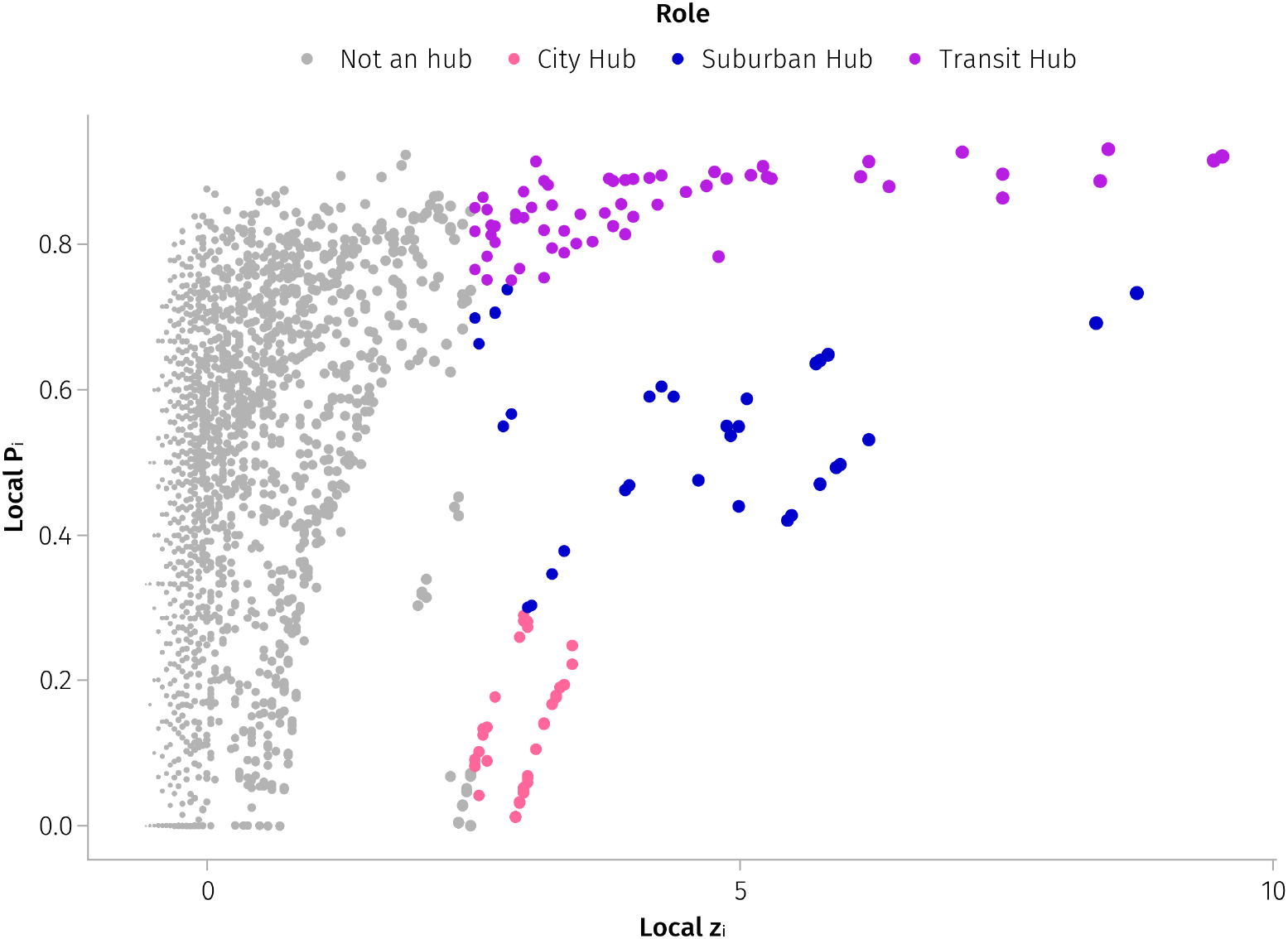
Local network cartography of the nodes in the PPI network. Each point is a node in the network, the point size is proportional to the node log degree. The local *z_i_* score quantifies normalised degree, and the local *P_i_* coefficient quantifies the relative density of connection for a node neighborhood.

The edges connected to the 184 hubs include over half of all the edges in the network, and the set of hubs and their first neighbours includes almost half of all genes in the network (2220).

We found that the genes in the network are significantly closer to the set of all hubs than random, as see in fig. 4 (A) there are significantly more nodes one step away from the set and significantly less nodes two steps away than in a random selection of a same sized set of nodes. This was accentuated for the set of the 61 Transit hubs, whilst the set of City and Suburban hubs tend to be further away from the rest of the networks, as visible in fig. 4 (B).

**Figure 4.**
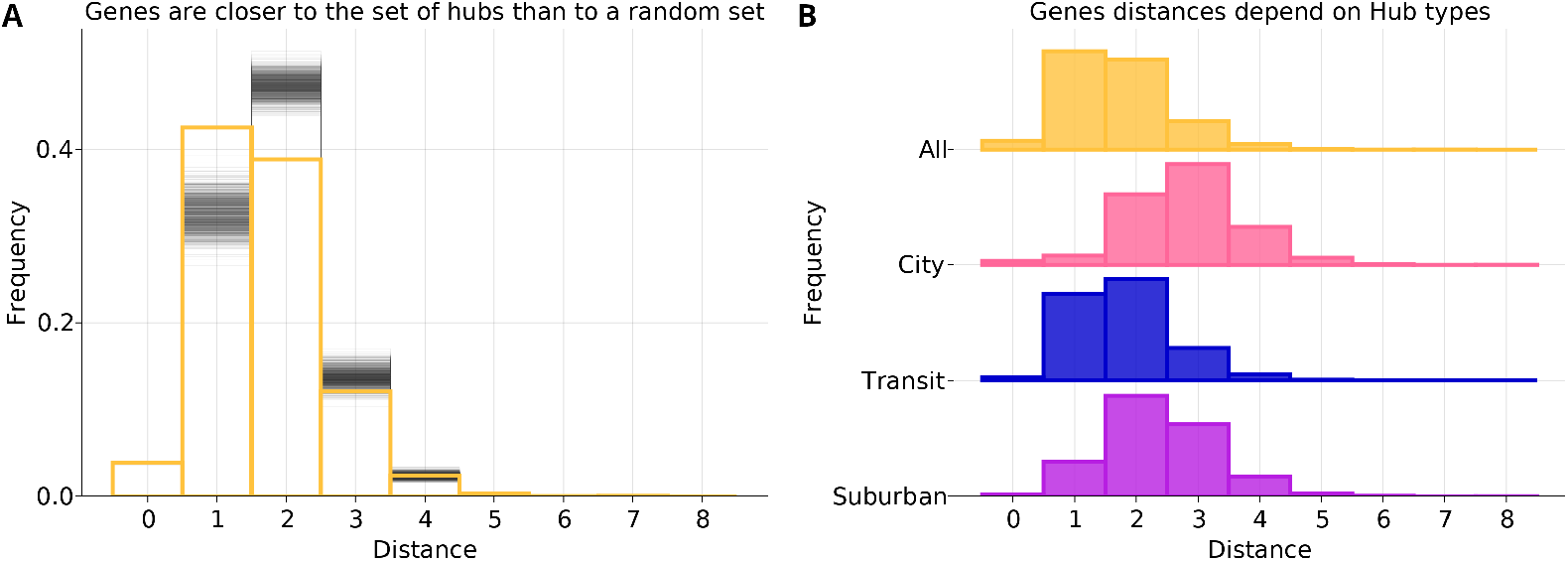
**(A)** the distribution of minimum distances from each node in the network to the set of all hubs; in yellow the distribution for the 184 hubs, in gray the distribution for each of the random replicates of 184 genes. **(B)** the distribution of minimum distances from each node in the network to each of the sets of specific hub types.

Potentially downregulated genes are predominantly Transit hubs (43%), whilst Transit hubs are 28% of the potentially upregulated genes. The proportions are reversed for City hubs (52% of potentially upregulated and 32% of potentially downregulated genes).

### 3.3. Gene Ontology enrichment

Analysing the 184 hub genes, we identified 341 GO biological process (GObp) significantly enriched. Of these, 131 terms are associated with more than 50 genes. Briefly, the terms of note include ones related to lipopolysaccharide (LPS), transforming growth factor *β* (*TGFβ*), cytokine production, reactive oxygen species (ROS), circadian rhythms, gene expression, G-protein coupled receptor (GPCRs), dopaminergic signalling, inflammatory response, and MAPK cascade.

We found 82 were significant terms for the City hubs, 24 terms for the suburban hubs, and 386 terms for the Transit hubs.

Dopamine is the only specific chemical signalling molecule appearing for both City and Suburban hubs. GPCR, phospholipase C, adenylate cyclase, and cytosolic calcium ion related terms also appear in both lists. RNA and viral processes appear in both Transit and Suburban hubs GO terms. LPS, circadian rhythms, inflammatory response, dopaminergic processes, and MAPK related signalling are common themes in the City and Transit hubs lists. The only chemical to appear in all 3 lists is dopamine. Dopamine appears in 5 terms in the City hubs list, and 1 term each in suburban and Transit hubs lists. No theme runs through all 3 type of hubs.

#### 3.3.1 City hubs

The terms enriched in City hubs are mainly related to signalling processes (44 terms). Of these, 16 are related to GPCRs, 9 of which are related to adenylate cyclase modulating receptors. We found 11 terms related to monoaminergic processes (5 related to dopaminergic processes). Glutaminergic, GABAergic, acetylcholinergic, purinergic and opioidergic terms also feature in small numbers, once again mainly to do with GPCRs. Calcium signalling and regulation of cytosolic calcium ion gradient terms appear as well, with 6 terms. In regards to immune system processes, inflammatory response, lipopolysaccharide signalling, chemokine signalling, and positive regulation of inflammation terms are enriched. Monocyte, neutrophil, dendritic cell, and general, cell chemotaxis also appear in the list.

#### 3.3.2 Suburban hubs

The Suburban Hubs featured 4 terms related to GPCRs, including both activating and inhibiting GPCRs coupled to adenylate cyclase. cAMP biosynthesis, protein kinase A and phospholipase C activity also occur, along with RNA transcription processes. We also found terms related to dopamine receptor signalling pathway and to viral transcription.

#### 3.3.3 Transit hubs

Transit hubs had the most terms related to immune processes. We found 16 terms related to cytokine signalling, 4 to B cells and 5 terms to T cells respectively, 3 each related to macrophages, leukocytes generally, and LPS. Interestingly viral process, regulation of defense response to virus, and negative regulation of NF-kB are also enriched, as well as 6 terms related to the MAPK cascade. We found terms related to growth factor and hormonal signalling but not to GPCRs. In total, 53 GO Terms include the word “signal.” Also of note, 9 terms related to RNA processes and 7 included the word DNA, of which 11 terms related to transcription. Histone deacyelation also featured twice, as well as both positive and negative regulation of gene expression, and histone phosphorylation. Circadian rhythmic processes appeared, including circadian regulation of gene expression. Positive regulation of reactive oxygen species production, cellular response to reactive oxygen species, and cellular response to hydrogen peroxide also appeared in the list. Finally, the transit hubs also had multiple energy production or mitochondria-related terms, including negative regulation of mitochondrial membrane potential and negative regulation of glycolytic process.

We performed clustering analysis for the 131 terms enriched from the hubs and identified 20 different groups at a similarity index cut off (*si*) of 0.45 and 31 groups at *si* = 0.7. These can be seen in fig. 5. As seen in the figure, at both *si* levels, clusters have been associated predominantly to immune processes, and in lesser measure to dopamine was also selected by and GPCR signalling pathway.

**Figure 5.**
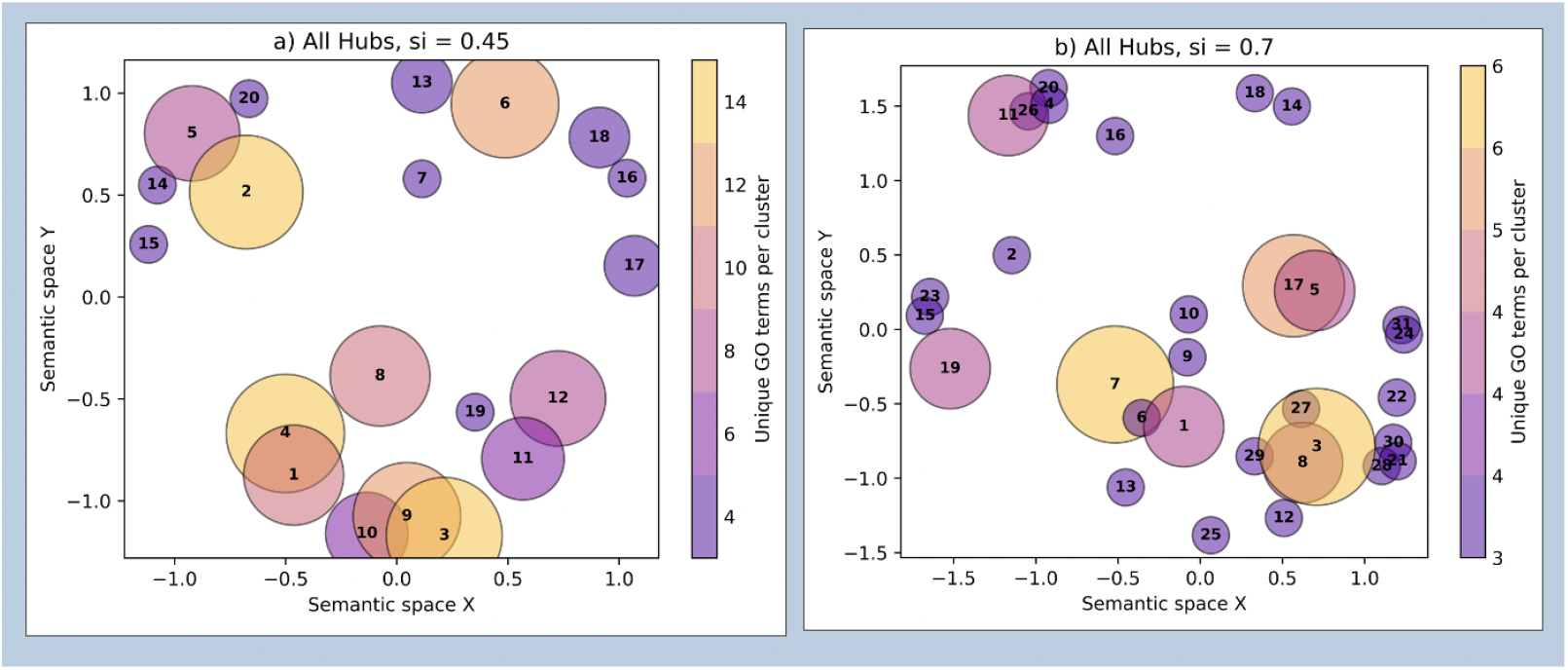
Clustering of GO Terms. (**a**): The clustering of 131 GO Terms by GO-Figure! at si = 0.45. Ordered by highest number of associated genes to lowest number of associated genes. Identity of the clusters in text. (**b**): clustering of the same terms by GO-Figure! at si = 0.7. Ordered by clusters with the highest number of associated genes to clusters with the lowest number of associated genes. Identity of the cluster in text.

In tbl. 2 we provide the identities of each cluster at the two *si* levels.

**Table 2.** Assigned themes for the GO terms cluster, at similarity index cut off of 0.45 and 0.7.

| cut off | Terms |
| --- | --- |
| $si = 0.45$ | 1. G protein-coupled receptor signaling pathway, 2. cellular response to dopamine, 3. positive regulation of cell population proliferation, 4. cytokine-mediated signaling pathway, 5. immune response, 6. sensory perception of pain, 7. cell-cell signaling, 8. positive regulation of cytosolic calcium ion concentration, 9. positive regulation of ERK1 and ERK2 cascade, 10. positive regulation of cell migration, 11. negative regulation of transcription by RNA polymerase II, 12. regulation of cytokine production involved in inflammatory response, 13. cell chemotaxis, 14. inflammatory response, 15. wound healing, 16. Aging, 17. locomotory behavior, 18. endocrine system development, 19. positive regulation of DNA-binding transcription factor activity, 20. response to hypoxia |
| $si = 0.7$ | 1. G protein-coupled receptor signaling pathway, 2. cell-cell signaling, 3. positive regulation of cell population proliferation, 4. cellular response to dopamine, 5. cytokine-mediated signaling pathway, 6. adenylate cyclase-activating G protein-coupled receptor signaling pathway, 7. regulation of cytokine production involved in inflammatory response, 8. positive regulation of apoptotic process, 9. negative regulation of transcription by RNA polymerase II 10. positive regulation of cytosolic calcium ion concentration, 11. response to lipopolysaccharide, 12. positive regulation of cell migration, 13. regulation of potassium ion transport, 14. leukocyte migration, 15. blood circulation, 16. wound healing, 17. ephrin receptor signaling pathway, 18. cell chemotaxis, 19. locomotory behavior, 20. response to cAMP, 21. positive regulation of ERK1 and ERK2 cascade, 22. regulation of small GTPase mediated signal transduction, 23. cognition, 24. Fe-gamma receptor signaling pathway involved in phagocytosis, 25. regulation of hormone secretion, 26. cellular response to reactive oxygen species, 27. positive regulation of cell growth, 28. positive regulation of protein kinase B signaling, 29. positive regulation of epithelial to mesenchymal transition, 30. positive regulation of phosphatidylinositol 3-kinase signaling, 31. stimulatory C-type lectin receptor signaling pathway. |

For the list of cluster members, see supplementary content.

Clustering of the individual hub types lists revealed a distinct structure to the terms annotated to each type, see fig. 6. The City hubs reduced to 14 clusters, with very little overlap. Manual analysis of the terms in each cluster revealed sets of processes which are related but may be better represented by a general term rather than the terms chosen by GO-Figure!. However, the terms themselves clustered into immune, signal transduction and chemical signalling processes. The suburban hubs clustered into 3 clusters of little relation. Most crucially, the Transit hubs clustered into 46 individual clusters with a huge degree of overlap. This may suggest that the Transit hubs do indeed represent genes involved in many biological processes which are not necessarily highly related to each other. Overall we found that the GO Terms resultant from the City hubs are more closely related than those of the Transit hubs.

**Figure 6.**
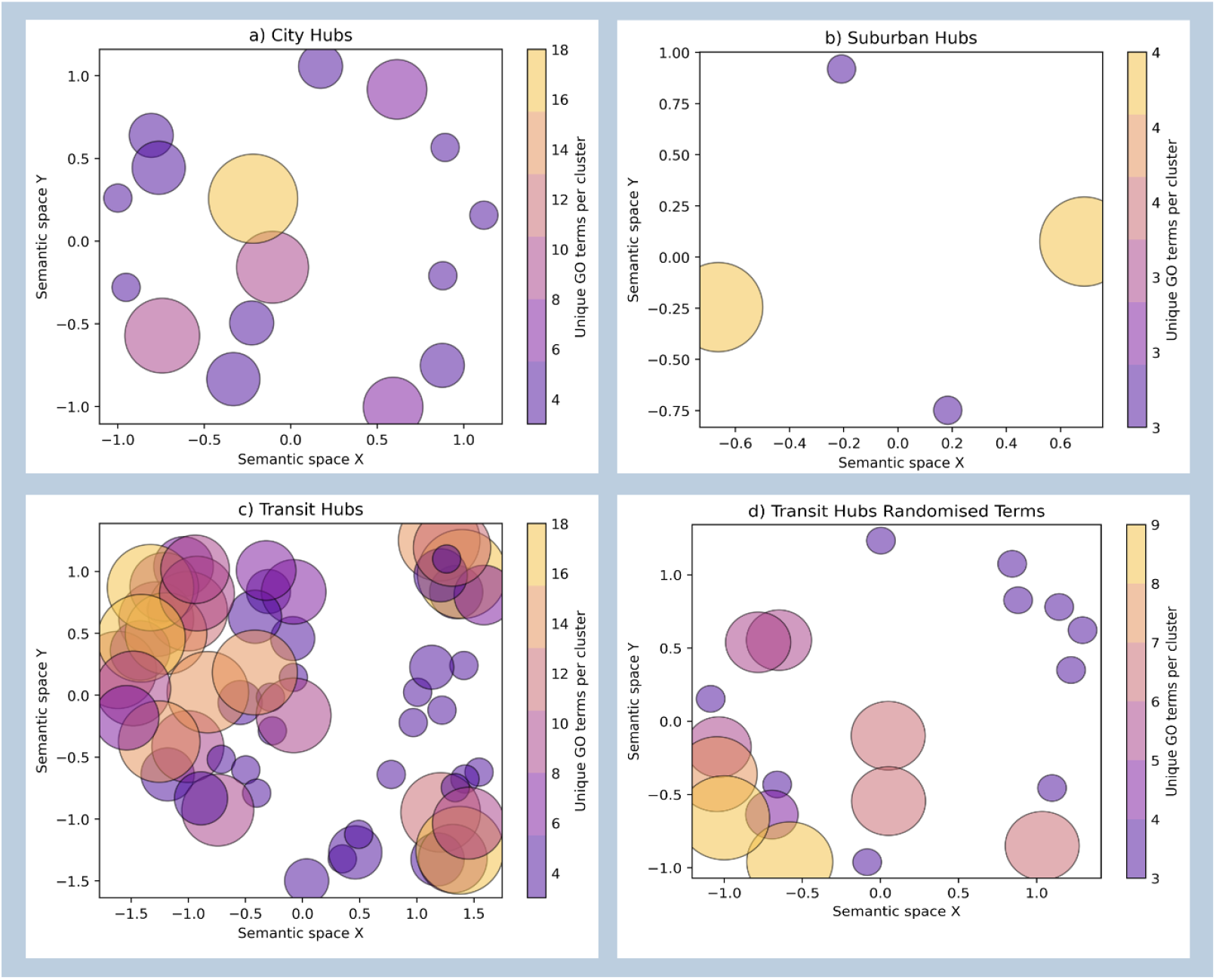
Clustering of GO Terms for Hubs. (**a**): The clustering of 82 GO terms enriched from the list of City hubs created on GO-Figure! (**b**): clustering of the 24 terms by GO-Figure! related to suburban hubs. (**c**): clustering of the 386 Transit hub GO terms. (**d**): clustering of 82 randomly selected Transit hub GO-terms for comparison to City hubs.

Validating our results against random replications, we found that for 197 of 200 Gene Ontology validation resulted in no statistically significant enriched GO terms at *α* = 0.05 (both for the set of all hubs and for each hub type hub set). All of the three analyses that had some significant GO terms had only one significant GO term in one analysis, therefore no clustering could be performed.

## 4 Discussion

For the first time we analysed the data from these three studies together, within a network analytic framework. The three studies we selected share the same platform, the same diagnostic criteria, and the same cell type. Thus, we considered the datasets highly compatible. The combined data includes 207 participants: almost double the size of the next largest methylation study of peripheral blood mononuclear cells (PBMCs) in ME/CFS (109 participants (Herrera et al., 2018)). This large sample size gives us more power to detect small logFCs. Two of the original studies (De Vega et al., 2017, 2014) identified 826 and 5,544 DMGs respectively. The other original study does not report how many genes were differentially methylated as it was looking for associations between methylation and single nucleotide polymorphisms (SNPs) (Herrera et al., 2018). Another study on methylation in ME/CFS found 17,296 differentially methylated probes, annotated to 6,368 DMGs (Trivedi et al., 2018). This suggests that whilst we did find 10,824 DMGs, it is not inconsistent with previous reports from smaller sample sizes. Hence, as we hypothesised, we found a large number of DMGs in ME/CFS PBMCs, often with a small LogFC.

### 4.1. Limitations

It is important to note that peripheral methylation does not necessarily represent methylation in the central nervous system (Non and Thayer, 2015; Walton et al., 2016). Because of the cell type analysed, the results discussed here focus on implications for the immune system. As we used publicly available data, we can not directly correlate the changes seen in methylation with symptomology or gene expression data of the same patient population. The PPI networks are intrinsically biased toward interactions that have already been studied. Likewise, the array platform used may be biased towards known cancer related genes as it was originally invented with these genes in mind (Barker et al., 2018; Non and Thayer, 2015). Thus, the hubs we identified may be biased in favour of proteins already well characterised and may be missing some less studied ones.

Moreover, whilst the gene expression can be indicated by the logFC methylation, the latter is not a perfect proxy: other factors may affect gene expression downstream of gene methylation. The relationship between non promoter methylation and gene expression is complex. Different studies produced contrasting results about the effect of gene body methylation, finding both positive and inverse relationships with gene expression, potentially depending on the methylation of other gene regions (Damgacioglu et al., 2019; De Almeida et al., 2019; Jjingo et al., 2012; Moore et al., 2013; Tang et al., 2017; Yang et al., 2014). Furthermore, the relationship may even be non-linear (Jjingo et al., 2012). While encouraging results have already been shown (Damgacioglu et al., 2019; Levy et al., 2020; Silva et al., 2021; Zhong et al., 2019), additional work needs to be done to robustly relate methylation changes with changes in gene expression. Moreover, histone aceylation and microRNA (miRNA) changes have been found in ME/CFS and these will change the transcription of genes (Almenar-Pérez et al., 2020; Ekua W. Brenu et al., 2014; Brenu et al., 2012; Jason et al., 2011; Petty et al., 2016).

### 4.2. Interpreting the set enriched terms

The results of the Gene Ontology enrichment analysis of the hub types pivot around four themes touching on wider epigenetic changes, immune system processes, signalling pathways, and dopaminergic process.

#### 4.2.1 Methylation changes underlying wider epigenetic changes

Our results strengthen the hypothesis that epigenetics may play a role in ME/CFS (Almenar-Pérez et al., 2019; Cortes Rivera et al., 2019; Vega et al., 2018). However, the methylation changes in ME/CFS may partially mediate the wider epigenetic changes found in ME/CFS. Considering the list of GO terms for transit hubs, we identified 6 abnormal pathways related to the regulation of miRNA production and transcription. Furthermore, Almenar-Pérez et al. (2020) found 8 upregulated miRNA in PBMCs of ME/CFS patients; our analysis independently predicts 5 of these genes would exhibit upregulated expression based on their methylation changes. Other studies have analysed miRNA expression in ME/CFS (Almenar-Pérez et al., 2020; Ekua W. Brenu et al., 2014; Brenu et al., 2012; Nepotchatykh et al., 2020; Petty et al., 2016) and one has attempted to integrate differential methylation with miRNA expression (Almenar-Pérez et al., 2019). We consider plausible that a degree of the miRNA abnormalities found in ME/CFS may be caused by differential methylation of their genes. Additionally, histone acetylation pathways also appear in the list of enriched terms from transit hubs. One study in ME/CFS patients found elevated levels of HDAC2 and HDAC3, suggestive of decreased gene transcription (Jason et al., 2011). Another suggests that hypoacetylation caused by upregulated expression of the HDAC family may contribute to post exertional malaise (McGregor et al., 2019), a key feature of ME/CFS symptomology (*Beyond myalgic encephalomyelitis/chronic fatigue syndrome: Redefining an illness.*, 2015; Fukuda et al., 1994). We found methylation changes suggestive of upregulation of multiple HDAC genes, including HDAC3. Thus, the widespread methylation changes we found may contribute to the other epigenetic abnormalities previously reported.

#### 4.2.2 Responses to perturbed immune system processes

In terms of immune system processes, a few GO terms stood out. Response to LPS and LPS mediated signalling appeared in the list of GO terms for both transit and city hubs. In fact, increased antibodies to LPS (Maes et al., 2012, 2007) and elevated blood levels of LPS (Giloteaux et al., 2016) have been reported in ME/CFS: this has been postulated to contribute to immune activation both centrally and peripherally (Morris and Maes, 2013). Another key molecule is reactive oxygen species (ROS). Terms related to response to ROS appeared in the transit hubs list, as well as positive regulation of their metabolism. ROS are suggested to contribute to ME/CFS as they are a crucial aspect of oxidative stress (OS) (Cortes Rivera et al., 2019; Jammes et al., 2012, 2005; A. Komaroff and Cho, 2011; Morris et al., 2019). OS related damage (DAMPs) may initiate and maintain an immune response whilst ROS may also disrupt mitochondrial functioning leading to reduced ATP production (Cortes Rivera et al., 2019; A. L. Komaroff and Cho, 2011; Morris and Maes, 2013). Unsurprisingly, we also found terms related to cytokine response. Finally, terms related to viral response appeared in the transit and suburban hubs enrichment. This is important because much work has been done exploring the association between ME/CFS and viruses (Bansal et al., 2012; Rasa et al., 2018). If viral responses are dysregulated by methylation changes, patients with ME/CFS may react differently to viruses than a healthy control would: relatively small changes to levels of O&NS, virus or LPS could be creating significant immune responses in an unpredictable manner.

Additionally, the GO terms relating to mitochondrial membrane potential and glycolytic activity are intriguing. There have been some findings of lowered mitochondrial membrane potential in patients with ME/CFS (Mandarano et al., 2020; Missailidis et al., 2020a, 2020b) as well as lowered glycolytic activities (Mandarano et al., 2020; Nguyen et al., 2019; Tomas et al., 2020), though the latter has been less consistent (Missailidis et al., 2021).

#### 4.2.3 Disruption of signalling pathways

Many GO terms were related to signalling processes. As discussed in the results section, the city hubs and suburban hubs had a high degree of GPCR associated processes enriched. These included a wide range of monoaminergic processes, including 5 dopaminergic processes. This may suggest that signalling by monoamines contributes to immune dysregulation in ME/CFS. Monoaminergic contribution to ME/CFS is not new, we present here evidence for a direct implication of monoaminergic signalling in immune dysfunction. The finding of dysregulated acetyl-cholinergic and opioidergic signalling also supports suggestions that these chemicals may contribute to ME/CFS immune dysregulation (Anderson and Maes, 2020; Marshall-Gradisnik et al., 2016; Tanaka et al., 2003; Wirth and Scheibenbogen, 2020). The same can be said of adrenergic signalling (Johnston et al., 2016; Loebel et al., 2016; Scheibenbogen et al., 2018; White et al., 2012). Calcium signalling also appeared in the GO terms list: it has been suggested that abnormalities in calcium signalling may contribute to natural killer (NK) cell hypofunction in ME/CFS (Cabanas et al., 2019; Eaton-Fitch et al., 2019; Nguyen et al., 2017). Considering the widely reported presence of autoantibodies to GPCRs in ME/CFS (Loebel et al., 2016; Tanaka et al., 2003; Wirth and Scheibenbogen, 2020), it is likely that methylation changes compound the effect of already disrupted GPCRs. In the transit hubs, terms related to immune signalling were present. Specifically, signalling mediated by both B and T cell receptors, cytokines generally, IL-6, TGF-b, and LPS. This may suggest dysregulated response to antigens and cytokines. As TGF-b was found to be the only consistently upregulated cytokine in one systematic review (Corbitt et al., 2019), it seems likely that TGF-b may underlie some of the immune dysregulation if its signalling mediation is also disrupted. In sum, signalling by neurotransmitters, immune receptors, and cytokines may be abnormal and may contribute to the immune pathophysiology of ME/CFS.

#### 4.2.4 Dopamine

Dopaminergic processes were the only processes to feature in the results for the GO enrichment analysis of all hub types. The fatigue in ME/CFS seems likely to be partially dopaminergic centrally, considering the efficacy of multiple dopaminergic agents in ME/CFS (Blockmans et al., 2006; Blockmans and Persoons, 2016; Crosby et al., 2021; Goodnick et al., 1992; Kaiser, 2015), and in mouse models of ME/CFS (Song et al., 2021; Thakur et al., 2020). This would also fit with the high rates of Attention Deficit Hyperactivity Disorder (ADHD) in ME/CFS (Sáez-Francàs et al., 2012), and monoamine metabolite levels in cerebrospinal fluid (Demitrack et al., 1992). Moreover, levels of both general and mental fatigue in ME/CFS have been inversely correlated with basal ganglia activation as assessed by fMRI, potentially secondary to dopaminergic deficits (Miller et al., 2014). Numerous other neuroimaging studies also support this notion (see (Almutairi et al., 2020; Cook et al., 2017; Gay et al., 2016; Josev et al., 2019; Manca et al., 2021; Shan et al., 2020; Shan et al., 2018; Tanaka et al., 2006; Van Der Schaaf et al., 2018, 2017; Wortinger et al., 2017; Wortinger et al., 2016)). Furthermore, Carandini et al. (2021) found an inverse correlation between levels of mental fatigue and DA tract abnormalities in MS. Carandini et al. (2021) suggested that this is evidence of dopaminergic abnormalities underlying mental fatigue, as hypothesised by Dobryakova et al. (2015). Research in animals, and in patients given cytokine therapies, has demonstrated that peripheral inflammation can lead to central dopaminergic hypofunction (Anisman et al., 1996; Felger et al., 2013; Felger and Miller, 2012; Lee et al., 2021). Hence it can be postulated that the immune dysregulation and inflammation in the periphery may lead to central dopaminergic deficits, which then may lead to some of the experience of fatigue in ME/CFS. It is important to note that this does not dismiss the impact of other mediators, such as impaired mitochondrial functioning (Holden et al., 2020; Mandarano et al., 2020; Tomas et al., 2017). Dopaminergic hypo-functioning would simply make the fatigue caused by peripheral processes feel significantly worse for the patients involved.

Our results suggest that dopaminergic signalling may also be abnormal in an immune context. The potential discovery of a dopaminergic element to ME/CFS immune dysregulation is consistent with its involvement in other autoimmune diseases including multiple sclerosis (Pacheco et al., 2014; Vidal and Pacheco, 2020a, 2020b). The effects of dopaminergic signalling in the immune system are wide, varied, and beyond the scope of this article. For review see (Arreola et al., 2016; Castorina et al., 2020; Feng and Lu, 2021). However, it is worth noting that signalling through the dopaminergic receptors D2 and D3, both of which are associated to hubs and potentially hyper-expressed, may have relevant immune effects. D2-like receptor agonists have been shown to bias T cells towards a Th2 type differentiation (Huang et al., 2010). Furthermore, D2 receptor activation is thought to inhibit natural killer cell activation and cytotoxicity (Capellino et al., 2020; Zhao et al., 2013). Of note considering the consistently found decreased NK cell cytotoxicity (Brenu et al., 2011; Fletcher et al., 2010; Huth et al., 2016; Levine et al., 1998; Maher et al., 2005; Marshall-Gradisnik et al., 2016) as well as a shift towards a Th2 response (Brenu et al., 2011; Broderick et al., 2010; Skowera et al., 2004; Torres-Harding et al., 2008) in ME/CFS.

It is worth noticing that suggestive evidences propose a role for dopamine in Covid-19 pathophysiology (Attademo and Bernardini, 2021b; Berber and Doluca, 2021; Nataf, 2020b). However, dopaminergic antagonists in ME/CFS would may have sedating effect. Hence, dopaminergic agents in ME/CFS may constitute the object of careful study, considering immune parameters, symptom perception, and side effects.

### 4.3. Different hubs play a different biological role

The participation coefficient is a combination of the clustering coefficient (Chalancon et al., 2013; Wasserman and Faust, 1994; Watts and Strogatz, 1998) and the original participation coefficient defined by Guimerà and Amaral (2005). However, whilst the clustering coefficient can be used to find modules of a network (Chalancon et al., 2013; Li et al., 2009; Nascimento, 2014; Zaki et al., 2013), and Guimerà and Amaral (2005)’s participation coefficient is based on modules, we use a combination of the two and a local averaging over network neighbours to overcome modularisation of an incomplete network. Our adapted participation coefficient highlights how much interaction there is between a group of neighbours: if a node has a high participation coefficient, its neighbours are sparsely connected, and the gene in question participates in many distinct biological processes. This is the case for transit hubs which GO enrichment terms are more numerous and cluster into smaller, more overlapping, clusters than either the terms for city hubs or suburban hubs, as we can see in fig. 6. The genes classified as transit hubs are commonly involved in signal transduction. Suggestively, whilst a few cytokine genes also appeared, transit hubs were associated to many immunity response terms.

City hubs, *P*_*i*_ < 0.3, are located in densely connected neighbourhood: therefore, their associated genes take part in well connected processes. By nature of their participation coefficient, they form functional units to a certain extent, which may interact with other units to a lesser degree. Hence an abnormality in a city hub may result in larger functional impact due to multiplying cascading effects (Shin et al., 2014), although this may be regulated by interaction strengths (Gaiarsa and Guimarães, 2019; Guimarães et al., 2018). From a biological perspective, this may suggest utility of drugs that modulate the signalling pathways seen from the GO enrichment. This can be seen in their clustering, where the clusters overlap less and contain more terms than the transit hubs.

The neighbours of suburban hubs may have varying degrees of connectivity between and within town groups of neighbours. However, whilst these hubs and their neighbours are not as interconnected as city hubs, there is still some degree of connectivity between their neighbours. These hubs almost seem to be the “go-betweens” of the city and transit hubs, as the terms enriched for them include both GPCR signalling and viral processes, not contained by the transit hubs and city hubs respectively.

### 4.4. Conclusions

Our result suggest that small, widespread, changes in gene methylation may mediate some of the immune system pathology seen in ME/CFS. If Covid-19 does indeed lead to ME/CFS as Komaroff et al. (2021) are suggesting, then methylation profiling a large sample of patients would be beneficial in understanding its pathogenesis. Efforts to understand the cause of methylation changes in ME/CFS, and Long Covid if applicable, should also occur. Moreover, analysing relationships between the DMGs is necessary: few signalling pathways that activate key effectors, their receptors are probably present in the hubs of this study. If these proteins do exist, they may lead to the discovery of druggable targets for ME/CFS immune dysregulation.

Our results further support the presence of methylation changes contributing to the immune dysregulation that characterises ME/CFS. They also suggest that responses to LPS, cytokines, O&NS, and viruses function differently to what is seen in healthy controls, exacerbating immune abnormalities beyond what would be expected otherwise. Finally, signalling by multiple endogenous chemicals is also perturbed and may affect immune activation. Depending on whether Long Covid is related to ME/CFS, our findings may be of interest for consideration when studying treatment implications.

